# A Sacrificial 3D Printed Vessel-on-Chip Demonstrates a Versatile Approach to Model Connective Tissue Pathology

**DOI:** 10.1101/2025.02.06.636821

**Authors:** Jonas Jäger, Phil Berger, Andrew I. Morrison, Hendrik Erfurth, Maria Thon, Eva-Maria Dehne, Susan Gibbs, Jasper J. Koning

## Abstract

For *in vitro* organ models, perfused vasculature is crucial to overcome nutrient diffusion limits and to generate immunocompetent models by allowing trans-endothelial migration of immune cells in and out of the tissue. However, vasculature is often disregarded due to its complexity to generate and the necessity to integrate flow. The aim here was to overcome these limitations by combining 3D printing and multi-organ-chip technology to generate a vascularized, fibroblast-populated connective tissue matrix on-chip. A 3D printed, sacrificial, water-dissolvable structure was incorporated into a multi-organ-chip to generate hollow channels within a collagen/fibrin hydrogel. Subsequently, the channels were populated with endothelial cells. Different hydrogel concentrations of fibrin were used to mimic healthy and early granulation tissue. The vessels were perfused, and stable metabolic/viability conditions (lactate dehydrogenase, glucose, lactate) acquired after 3 days for 7 days total. In high fibrin gels, angiogenic sprouting and increased secretion of angiogenic cytokines was observed. Perfusion with monocytes revealed differentiation into macrophages and migration across the endothelium into the tissue. In conclusion, the versatile, easy method to pattern hydrogels in multi-organ-chips can serve as the basis to build the next generation of vascularized, immunocompetent human organ models, and opens new possibilities to study health and disease.

## 1. Introduction

In the human body, all cells inhabit a three-dimensional (3D) tissue-specific niche. They are embedded in a network of acellular molecules called the extracellular matrix (ECM), which provides structural support and biochemical cues. Molecular composition is highly variable and can change with age, tissue damage and disease (tumor).^1^ In order to maintain homeostasis, cells require a consistent supply of oxygen, nutrients, as well as the removal of waste products. To facilitate this, the ECM is penetrated by the blood vasculature, which comprises a complex system of arteries, veins, and capillaries. Furthermore, the systemic circulation enables communication across organs through factors such as soluble mediators, exosomes or cells to maintain viability and homeostasis.^2, 3^ The inner layer of blood vessels and capillaries is directly in contact with the circulating blood and consists of a single layer of endothelial cells (ECs), known as the endothelium. Maintaining its integrity, ECs are connected via tight junctions and adherens junctions, transmembrane proteins such as platelet endothelial cell adhesion molecule 1 (PECAM-1; also known as CD31) and vascular endothelial cadherin (VE-cadherin; also known as CD144) to form a barrier.^4^ However, cell-cell junctions are not always tightly closed. At sites of infection or injury, coordinated disassembly of junction proteins leads to the extravasation of immune cells such as monocytes, neutrophils or lymphocytes at specific hotspots out of the blood into the surrounding tissue to elicit an immune response.^5, 6^ Predominantly, but not exclusively, during development, blood vessels are formed either by vasculogenesis, *de novo* from endothelial progenitors, or by sprouting from pre-existing ones in a process known as angiogenesis.^7^ Vascular sprouting is also common in the fibrin-rich granulation tissue that forms in wound beds. This can lead to the formation of scars (fibrosis).^8^

In recent years, engineering of human three-dimensional (3D) *in vitro* models has improved significantly.^9^ Advanced models are capable of recapitulating human biology and creating more physiologically relevant human organ and tissue models. Microphysiological systems (MPS) have emerged, which are models mimicking human tissue and organ function *in vitro* at the micrometer scale. A type of MPS are Organ-on-Chip (OoC) systems which allow continuous organ, organoid or tissue model perfusion with very small fluid volumes.^10, 11^ Multi-organ-chips (MOCs) take this OoC concept further, emulating human complexity and allowing crosstalk between different organ models. This enables drug testing and disease modelling on a systemic level, which is particularly important when studying drug absorption, distribution, metabolism, and excretion (ADME) as well as modelling inflammation and immunological diseases.^3, 12^

Modeling perfused blood vasculature presents a challenge, as it comprises the combination of an OoC with ECs, within an ECM to allow angiogenic sprouting for disease models and the integration of other relevant cell types such as fibroblasts, pericytes and smooth muscle cells. Several perfused vascularized organ models have been generated already.^13–20^ However, the majority of these models rely on gravity-driven or laminar, non-recirculating flow with soft tubing which complicates multi-organ cultures. In addition, these studies utilize highly customized in-house fabricated chip setups, which are not commercially available and are very challenging to reproduce. Furthermore, several barrier models of e.g. skin, gut and lung make use of a transwell system on which the epithelium is grown on the upper side of a porous membrane with perfusion on the lower side.^21–23^ This configuration makes it almost impossible to generate a perfused vascular network within the barrier model.

The aim of this study was to generate and characterize a vascularized, perfused fibroblast-containing connective tissue hydrogel directly inside a MOC platform and perfuse it with peripheral blood-derived human monocytes. The TissUse HUMMIC chip was used with micropumps directly on-chip, driven by a control unit which applies pressure and vacuum to generate perfusion. As OoC platforms are required to mimic healthy and diseased tissues, the objective was to highlight the versatility of the platform by modifying the hydrogel composition. Both collagen and fibrinogen are widely used for constructing hydrogels in tissue engineering. However, fibrin is also involved in granulation tissue formation, blood clotting and the inflammatory response of early wound healing which eventually can lead to fibrosis.^8, 24^ Accordingly, two different hydrogels were compared: a low fibrinogen (F_low_) hydrogel to mimic a healthy tissue and a high fibrinogen (F_high_) hydrogel to mimic early granulation tissue. The vessel within the hydrogel was perfused for up to 7 days, while metabolic markers and viability of the EC-fibroblast co-cultures were assessed throughout the experiment. Following perfusion of monocytes, the differentiation into macrophages and migration into the tissue was evaluated. Overall, perfusable blood vessels were generated on-chip and it was demonstrated that the hydrogel composition can be modified to mimic early granulation tissue.

## 2. Results

### 2.1. Manufacturing of the chip platform

The HUMIMIC Chip2 24-well (TissUse, Berlin, Germany) was utilized and produced as described earlier.^25^ In brief, a microscopic glass slide was attached to a polydimethylsiloxane (PDMS) layer incorporating a closed loop of microfluidic channels. The circulation can be accessed from the top via the two cell culture compartments, one with a diameter of 6.5 mm (vascular compartment) and one with a diameter of 14 mm (hydrogel compartment). The PDMS layer was attached to a polycarbonate adapter plate on the upper side, which served as a housing for cell culture compartments, lids, and pump adapters (**Figure 1A**). To apply flow, the on-chip micropumps were actuated by connecting the pump adapters via tubing to a HUMIMIC Starter control unit (TissUse). A 3D-printed sacrificial polyvinyl alcohol (PVA) structure was printed outside of the chip, incorporated in the hydrogel compartment, and aligned with the on-chip circulation when plasma-bonding the glass slide to the PDMS layer (**Figure 1A and Supplementary Figure S1A**). The versatility of 3D printing allowed for multiple vascular designs to be printed. However, in this study, either a straight line or a bifurcated print was incorporated into the chips (**Figure 1B**, i and ii).

**Figure 1:**
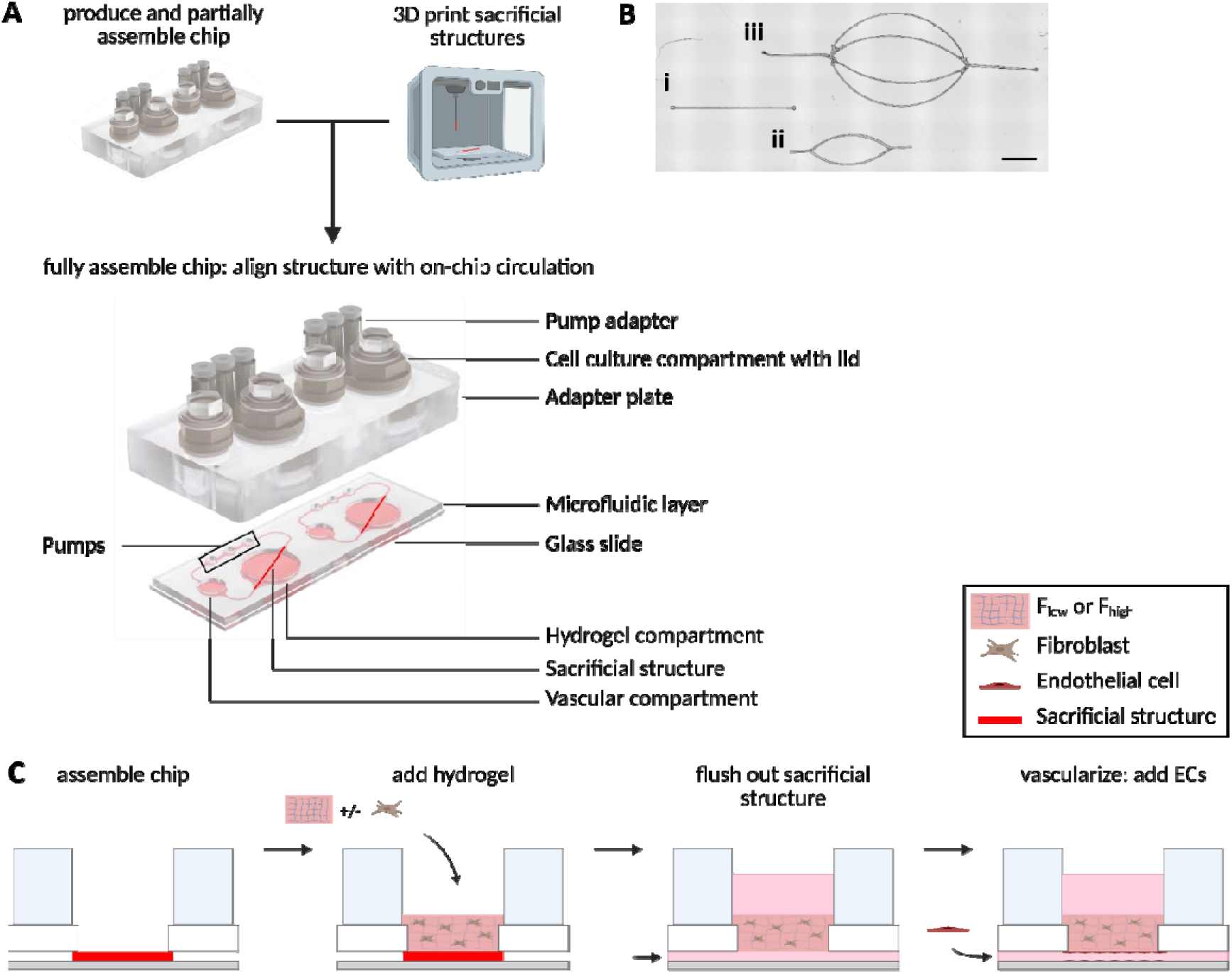
Sacrificial structure incorporated into the multi-organ-chip (MOC) platform to generate perfused vasculature. (A) Production of the chip by the integration of a 3D printed sacrificial structure in the MOC. The structure was incorporated in the hydrogel compartment and aligned with the on-chip circulation. One MOC consists of two independent circuits with a closed microfluidic loop. (B) 3D printing allows generation of different print designs. For simplification, design (i) and (ii) were used in this study. Scale bar: 3000 µm. (C) Cross-sectional view of the hydrogel compartment. Perfused vasculature was generated in three steps: addition of the hydrogel, removal of the sacrificial support structure and subsequent seeding of endothelial cells. F_high_: high fibrinogen, F_low_: low fibrinogen hydrogel concentrations.

### 2.2. Hydrogel channel generation and characterization of flow profiles and shear stress

After manufacturing of the MOC, channels were generated within the hydrogel directly on the chip. Hereto, a hydrogel was layered on top of the sacrificial structure. While the PVA structure was dissolved, the hydrogel polymerized which resulted in a hollow channel within the hydrogel, connected to the on-chip microcirculation (**Figure 1C**). Since it was unclear how the introduction of a hydrogel channel would change the flow within the chip, flow profiles and shear stress were first evaluated. Before the incorporation of cells, flow and shear stress in the hydrogel vessel were characterized and at distinct locations in the on-chip microcirculation (**Figure 2A**).

**Figure 2:**
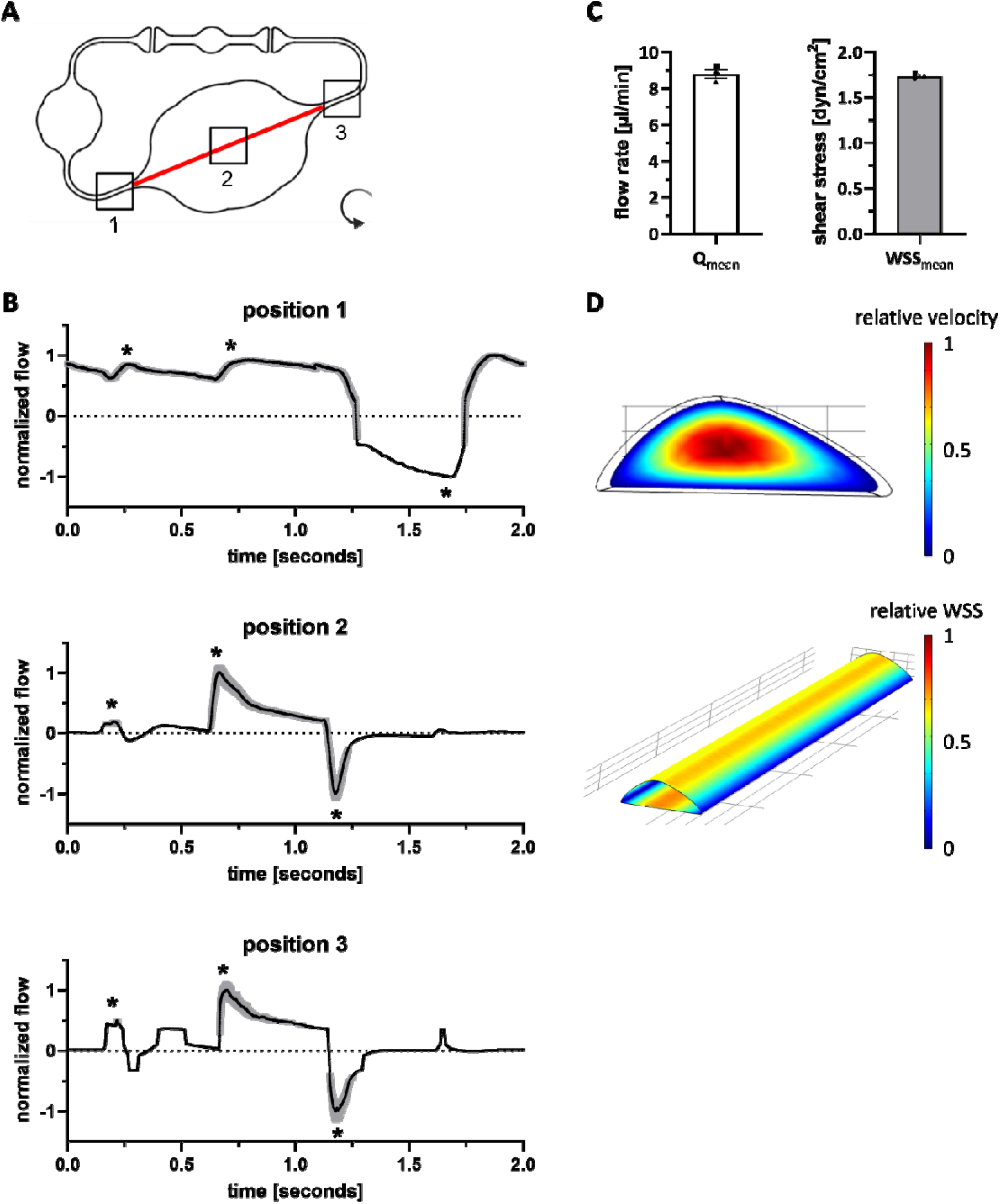
Hydrogel channel characterization: flow rate and shear stress. (A) Microfluidic profile of the modified Chip2-24. Non-invasive micro-particle image velocimetry (µPIV) was measured at various positions (1,2 and 3), as indicated by the black squares. Arrow indicates the direction of perfusion. (B) Normalized flow profile at respective position in the chip over a time interval of 2 seconds. Shown is Q_mean_ in black and Q_std_ in gray. Asterisks indicate extrema (2 peaks followed by a backflow). Pump settings: anti-clockwise, ± 500 mbar, 0.5 Hz. (C) Mean flow rate (Q) and wall shear stress (WSS) on-chip. Q and WSS were calculated as average from position 1 and 3. Mean ± SEM of n = 3 individual microfluidic circuits is shown, represented as different symbols. (D) Spatial distribution of velocity in the cross section and WSS in a section of the hydrogel channel.

Notably, at all positions in the chip (before, in and after the hydrogel vessel) a pulsed, pump-characteristic flow profile with two peaks followed by a short backflow was observed (**Figure 2B**).^26^ Position 1 and 3 showed similar PIV and were used to calculate the mean flow rate and the mean wall shear stress (WSS) as the channel geometry was well defined in the PDMS channels. The mean flow rate in the channels was 8.8 ± 0.24 µl/min and the mean WSS was 1.7 ± 0.018 dyn/cm^2^ (**Figure 2C**). Since the channel geometry in the hydrogel channel (position 2) was not well defined, a computational fluid dynamics simulation was performed to assess mean flow rates and WSS. This simulation shows the highest velocity in the center of the cross section of the hydrogel channel and the highest WSS in the center of top and bottom of the semi-elliptical channel walls, 7-times higher compared to the sides of the channel (**Figure 2D**).

### 2.3. Generation of a functional vessel by endothelial cell population

After flow rate and shear stress characterization, a blood vessel was generated within the hydrogel (**Figure 1C**; for timeline, see Supplementary Figure S1B). Hereto, the hollow channels of a bifurcated sacrificial print were populated with primary, dermal-derived ECs and the vessels cultured in flow for 7 days (**Figure 1B and 3A**). Notably, ECs not only populated the entire channels (sides, top and bottom), but also aligned with shear stress in the direction of flow within the hydrogel vessel (**Figure 3B**). A lumen was generated with a cross-section of semi-elliptical shape by the negative mold of the print (**Figure 3C** and Supplementary Movie S1). To assess barrier properties of the generated vessel, a FITC-dextran permeability assay was performed in a straight vessel. After 10 minutes, 70 kDa FITC-dextran was only detected strongly within the vessel, while no FITC-dextran was detected within the hydrogel. In contrast, channels without ECs showed clear diffusion of FITC-dextran into the hydrogel (**Figure 3D**). Overall, a functional vessel could be generated on-chip by population of the hydrogel channels with ECs.

**Figure 3:**
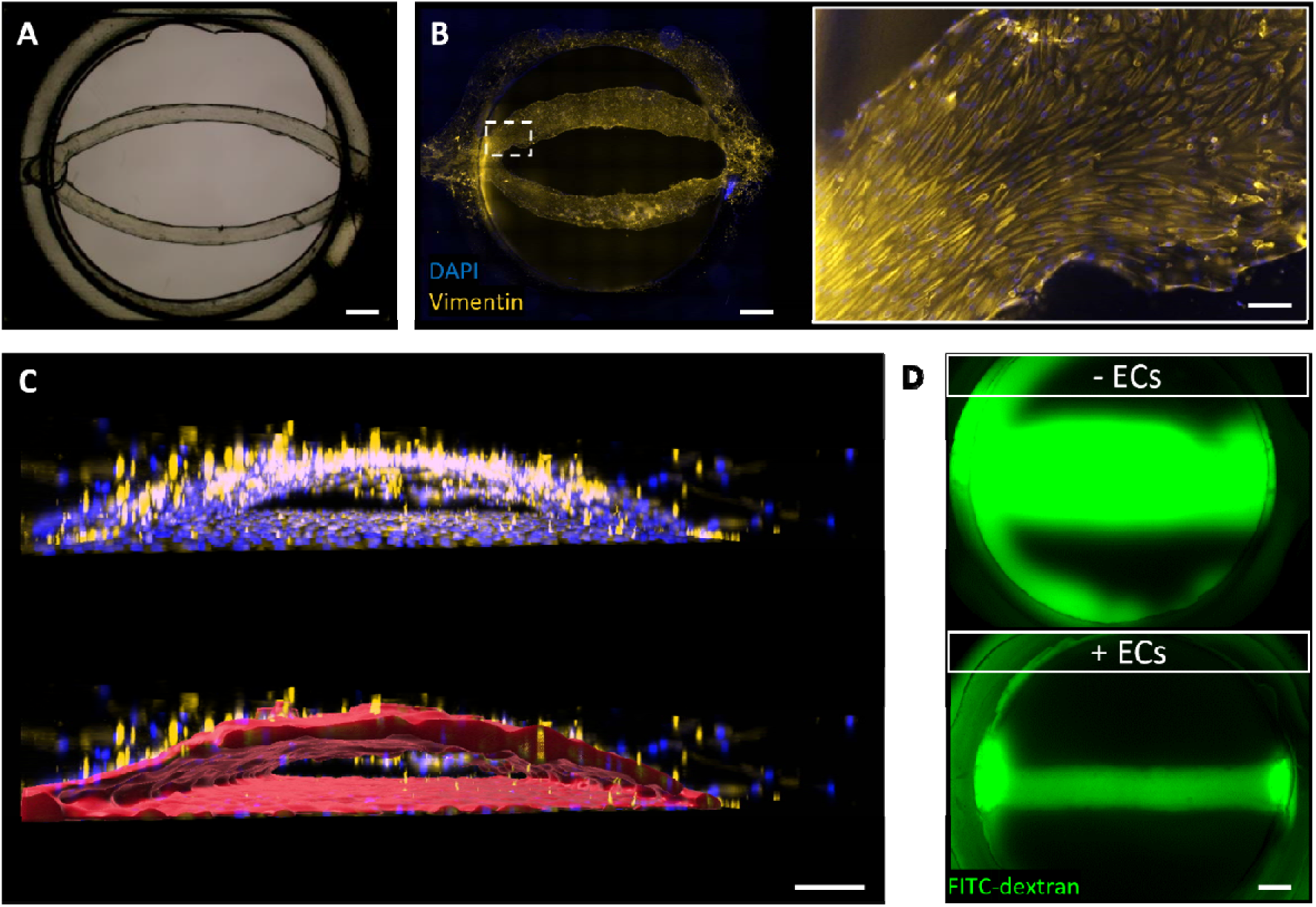
Endothelial cells grow on channel walls and form a functional, perfusable vessel on-chip. **(A)** Bright field image of a bifurcated sacrificial structure inside the hydrogel compartment. **(B)** Bottom view of the compartment at day 7. Endothelial cells (ECs) in chip populate the channel walls and align in direction of flow. **(C)** Surface rendering (red) visualizes hollow channels and semi-elliptical cross section. **(D)** FITC-dextran perfusion in channel with or without ECs shows perfusability and barrier properties of the blood vessel. ECs: Endothelial cells, FITC: Fluorescein isothiocyanate. PDMS: polydimethylsiloxane. Scale bars: 800 µm for overviews, 100 µm for magnification in (B) and 200 µm for magnifications in (C).

### 2.4. Hydrogel composition influences fibroblast and vessel morphology

Next, two hydrogel conditions were compared: low fibrinogen (F_low_) hydrogel to mimic the healthy state and high fibrinogen (F_high_) hydrogel to model early granulation tissue. In addition to the ECs in the channels, fibroblasts were incorporated into the hydrogel to model the connective tissue.

After 7 days, angiogenic sprouting was observed only in F_high_ hydrogels where ECs started to invade the fibrin matrix. This was not observed in F_low_ hydrogels where the vessel maintained the architecture without morphological changes (**Figure 4A** and **B**, i). In addition, morphology of the fibroblasts changed noticeably from star-shaped cells in F_high_ to elongated cells in F_low_ (**Figure 4B**, ii and iii). Channel width increased after addition of a hydrogel on top of the sacrificial structure, which had an initial width of 420 ± 3 µm after printing (**Figure 4C**). This widening was influenced by the composition of the hydrogel and was more pronounced in F_high_ hydrogels (605 ± 20 µm) than in F_low_ hydrogels (481 ± 6 µm). As shown by the spread of the data points, the variability of the channel width also increased with the addition of ECs and fibroblasts. Variability in hydrogel rheology between F_high_ and F_low_ hydrogels could have caused these differences in channel width and angiogenic sprouting. Therefore, the Young’s modulus of the F_high_ and F_low_ hydrogels was determined by nanoindentation. Notably, F_low_ hydrogels (95.03 ± 8.11 Pa) were 9x stiffer compared to F_high_ hydrogels (847.04 ± 80.22 Pa) (**Figure 4D**). Overall, these results emphasize that the hydrogel composition can influence angiogenesis, fibroblast morphology, vessel width and hydrogel stiffness.

**Figure 4:**
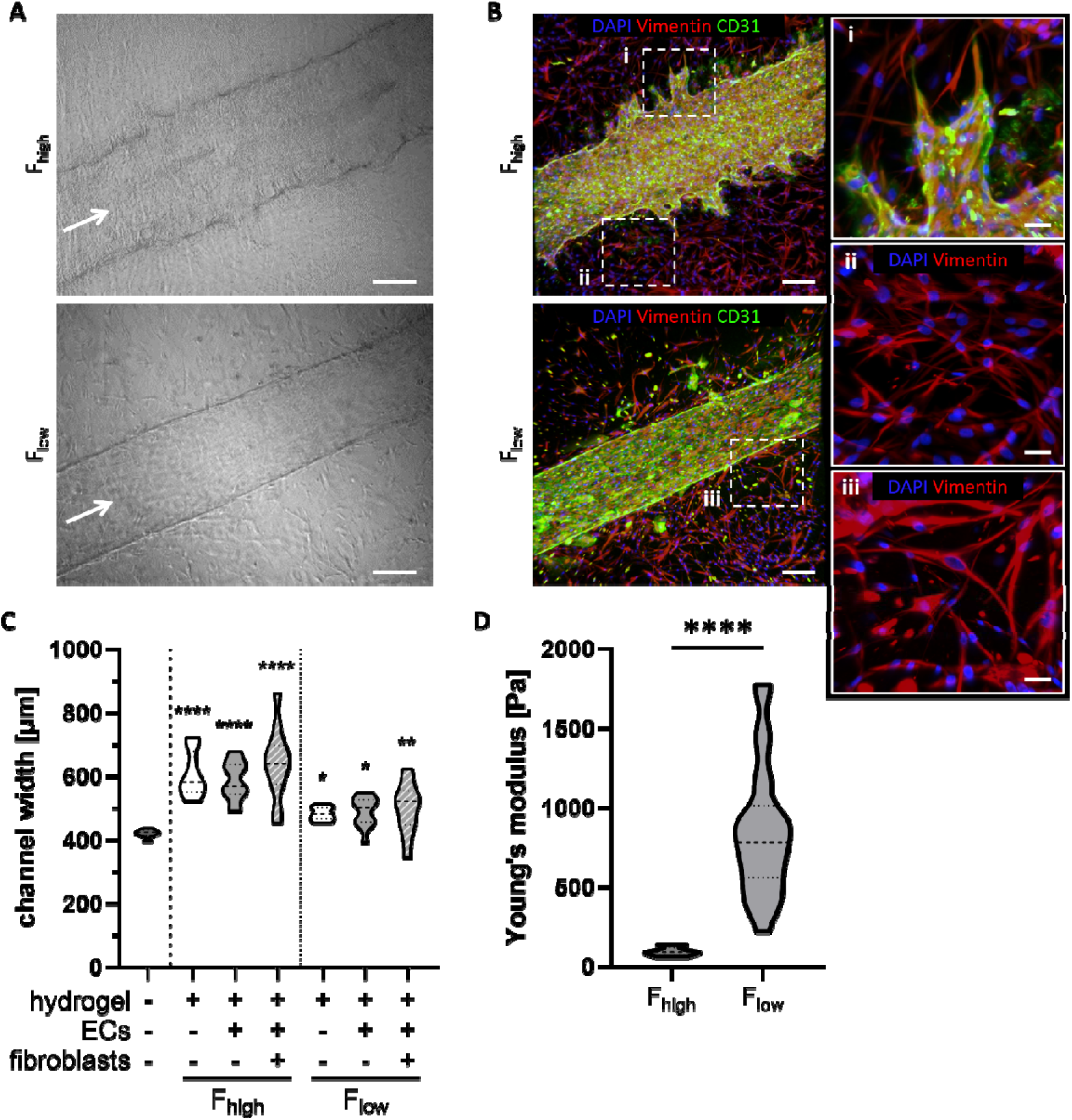
Hydrogel composition influences angiogenic sprouting, fibroblast morphology, channel width and stiffness. **(A)** Bright field images showing vessel morphology in F_high_ and F_low_ hydrogels. White arrow indicates direction of perfusion. Scale bars: 200 µm. **(B)** Immunofluorescent staining of fibroblasts (vimentin, red) and ECs (CD31, green) in F_high_ and F_low_ hydrogels. Magnifications visualize angiogenic sprouting (i) and fibroblast morphology (ii, iii) within respective hydrogels. Maximum intensity projections are shown. Scale bars: 200 µm in overviews, 50 µm in magnifications. All images were taken at day 7. **(C)** Width of the vessel within F_high_ and F_low_ hydrogels with and without ECs and/or fibroblasts. The sacrificial print without a hydrogel was used as a control. Each condition contains n ≥ 11 data points, which are compared to the control (-/-/-) with a one-way ANOVA. ECs: endothelial cells. **(D)** Young’s modulus of hydrogel conditions, measured after 4 days in the chip with fibroblasts and ECs. Each condition with n ≥ 10 data points, compared with an unpaired t-test. ****p ≤ 0.0001, **p ≤ 0.01, *p ≤ 0.05.

### 2.5. Pro-inflammatory cytokine secretion increased in hydrogel medium and higher angiogenic cytokine secretion in F_high_ hydrogel

As angiogenic sprouting was observed into F_high_ but not into F_low_ hydrogels, secretion of angiogenic (bFGF, VEGF, sPECAM-1, EGF, angiopoeitin-1 and PlGF) and pro-inflammatory cytokines (TNFα, IL-6, IL-8) was investigated. Since the chip setup allowed for the separate sampling of culture supernatant from the vascular and from the hydrogel compartment (**Figure 1**), this also enabled the comparison of cytokine secretion within the vasculature and on top of the hydrogel (**Figure 5A**).

**Figure 5:**
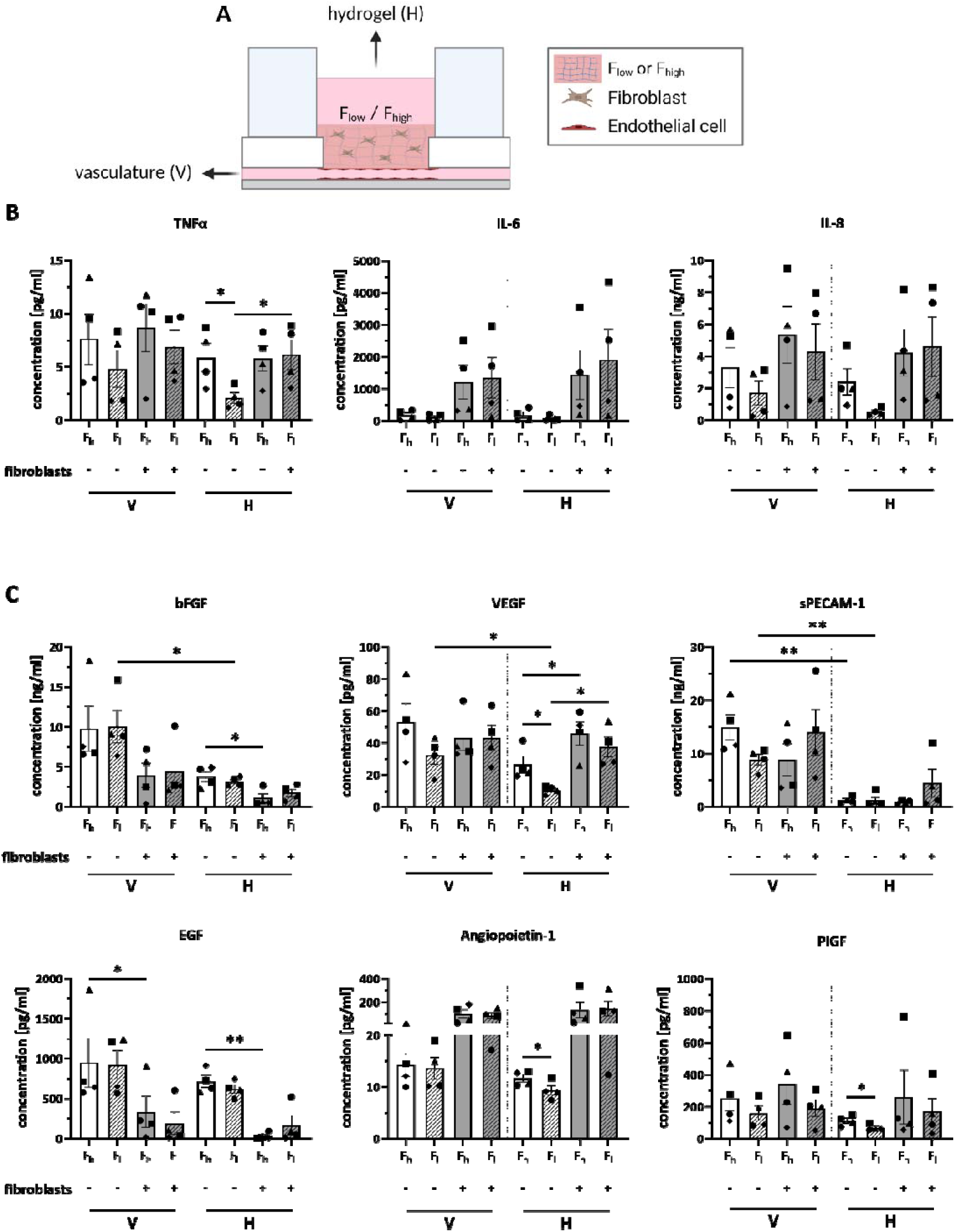
Elevated pro-inflammatory and angiogenic cytokine secretion in F_high_ hydrogel. **(A)** Medium was sampled either from the vasculature or the hydrogel at day 7. Hydrogel was added with or without fibroblasts in F_high_ and F_low_ hydrogels. **(B)** Pro-inflammatory cytokine and **(C)** Angiogenic cytokine concentrations. F_h_: high fibrinogen hydrogel, F_l_: low fibrinogen hydrogel; V: vasculature, H: hydrogel. Symbols represent different donors. Data is represented as mean ± SEM; n = 4 independent experiments performed with a minimum of two intra-experimental replicates; **p ≤ 0.01, *p ≤ 0.05, tested with a one-way ANOVA.

Within the hydrogel compartment, comparing F_high_ to F_low_ hydrogel conditions with ECs but without the addition of fibroblasts, revealed an increased secretion of TNFα, VEGF, angiopoietin-1 and PlGF into the F_high_ hydrogel compartment (**Figure 5B, C**). Interestingly, within the same hydrogel condition, the concentrations of TNFα (F_low_) and VEGF (F_low_ and F_high_) were higher when fibroblasts were present, while bFGF (F_high_) and EGF (F_high_) concentrations were lower. Also in the vascular compartment, a decrease in EGF was observed upon addition of fibroblasts to the F_high_ hydrogel (**Figure 5C**). When comparing between the vascular and the hydrogel compartment within the chip, higher secretion of bFGF, VEGF and sPECAM-1, into the vascular compartment was detected within the same hydrogel condition without fibroblasts (for bFGF and VEGF in F_low_ hydrogels only and for sPECAM-1 in F_low_ and F_high_ hydrogels)(**Figure 5C**).

In summary, although higher levels of VEGF, angiopoietin-1 and PlGF were observed in F_high_ compared to F_low_ hydrogels without fibroblasts, there were no clear differences in these angiogenic factors between F_high_ and F_low_ hydrogels that contain fibroblasts.

### 2.6. Vessel co-cultures are metabolically stable and viable in chips for 7 days

To determine whether the cells in the hydrogels were metabolically active and viable in the chip over time, glucose, lactate and LDH concentrations were measured in the culture medium. First, basal glucose levels of the vasculature and the hydrogel medium were assessed (indicated by dotted lines) and glucose levels in the whole circulation calculated (**Figure 6**, dashed line). In the vasculature compartment, glucose levels were lower when fibroblasts and ECs were present compared to ECs alone, indicating higher metabolic activity with the addition of fibroblasts. Interestingly, glucose levels in the vasculature were lower in F_high_ hydrogels when compared to F_low_ hydrogels (**Figure 6A**). In line with glucose measurements, lactate concentration was also higher when fibroblasts and ECs were present compared to EC alone. The highest lactate level was observed in the F_high_ hydrogel with fibroblasts at day 7 (**Figure 6B**).Except for glucose concentrations in the hydrogel compartment, glucose, and lactate concentrations in the vasculature and on top of the hydrogel increased across all conditions from day 0 to 3 and then remained stable until day 7. LDH release, an indication for cell death and membrane porosity, reached its peak in all conditions at day 3 and then decreased from day 5 onwards (**Figure 6C**). The increase in LDH levels was less pronounced in the hydrogel compartment compared to the vasculature compartment.

**Figure 6:**
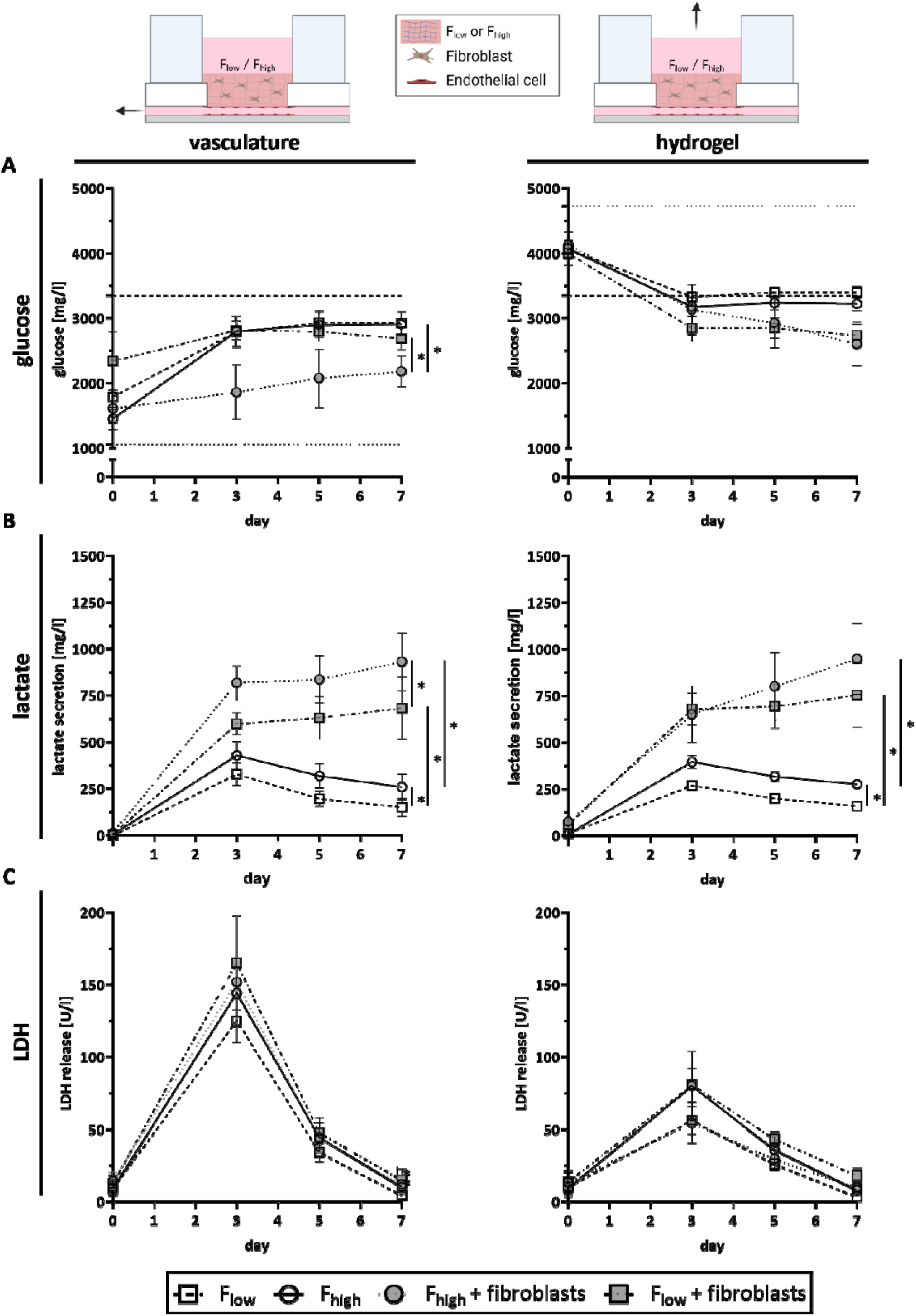
Co-cultures are metabolically stable and viable in the chip over time. Measurement of **(A)** glucose, **(B)** lactate and **(C)** Lactate dehydrogenase (LDH) throughout the 7 day co-culture period on-chip. Glucose and lactate were used as markers for cell metabolism and LDH as a cell death marker. Medium was sampled either from the vasculature compartment or on top of the hydrogel. Dotted lines indicate basal glucose levels of respective vasculature and hydrogel medium and dashed lines the basal glucose levels in the whole chip. Symbols represent different conditions as mean ± SEM, **p ≤ 0.01, *p ≤ 0.05, tested with one-way ANOVA at day 7. n = 4 independent experiments performed with a minimum of two intra-experimental replicates.

Taken together, these results show that the metabolic biomarkers glucose consumption and lactate secretion increased up to day 3 and stabilized over time until day 7. This coincides with LDH release which reached its maximum at day 3 and gradually decreased afterwards. Accordingly, a metabolically stable co-culture of ECs and fibroblasts is achieved in the chip after 3 days and remains stable until at least day 7.

### 2.7. Inter- and intra-experimental variation

To assess the reproducibility of the methodologies, we additionally determined the inter- and intra-experimental variability for glucose, lactate and LDH levels as well as cytokine secretion at day 7. Between individual experiments (inter-experimental) and within the replicates of a single experiment (intra-experimental), minor variation in glucose and lactate levels was observed in F_high_ and F_low_ cultures when only ECs were present (Supplementary Figure S2). This inter- and intra-experiment variation increased slightly when fibroblasts were incorporated into the hydrogel. For LDH release, a similar moderate level of inter- and intra-experimental variation was observed independent of the experimental condition.

When the cytokine secretion was very low, then the observed inter- and intra-experimental variation was low. This variation increased between different repeats and within individual experiments when higher cytokine secretion was observed (Supplementary Figure S3).

### 2.8. Circulation of monocytes

Finally, to mimic an inflammatory vascular setting, monocytes were circulated through the vessel and their differentiation into macrophages and migration into the hydrogel was assessed. Blood-derived primary CD14^+^ monocytes were added to the vascular compartment and circulated through the blood vessel for 24 hours before microscopy and flow cytometry analysis (**Figure 7A**). After addition of the monocytes, no significant change in LDH concentrations was observed (**Figure 7B**). However, LDH concentrations were generally slightly higher in the F_low_ hydrogel, especially within the vasculature (6.3 U/l vs. 15.1 U/l increase). Hydrogels were nanoindented directly in the chips to assess the rheology after the addition of immune cells. Similar to the hydrogels without monocytes, the hydrogel stiffness was higher in F_low_ than in F_high_ conditions (Supplementary Figure S4E). For flow cytometry analysis, single cell suspensions were harvested directly from the vasculature compartment and from the enzymatically digested hydrogel. In comparison to a static control, the majority of the monocytes on the chip expressed HLA-DR and CD68, indicating that they differentiated into a macrophage-like phenotype (**Figure 7C**). This was observed to be independent of the hydrogel composition. However, the mean fluorescent intensity (MFI) of CD68 was significantly elevated in F_high_ hydrogels, implying a more pronounced macrophage phenotype (**Figure 7D**). Further characterization revealed that the cells did not express CD40 (Supplementary Figure S4D) but the percentage CD206 expressing cells was increased throughout all conditions (**Figure 7E**), suggesting an M2-like phenotype. Although imaging data indicated that a lot of the immune cells attached to the endothelium (**Figure 7F**, i), a minor proportion migrated deeper into the hydrogel across the endothelial barrier (**Figure 7F**, ii). In summary, this shows monocyte-macrophage differentiation on-chip with and trans-endothelial migratory capacity of the immune cells.

**Figure 7:**
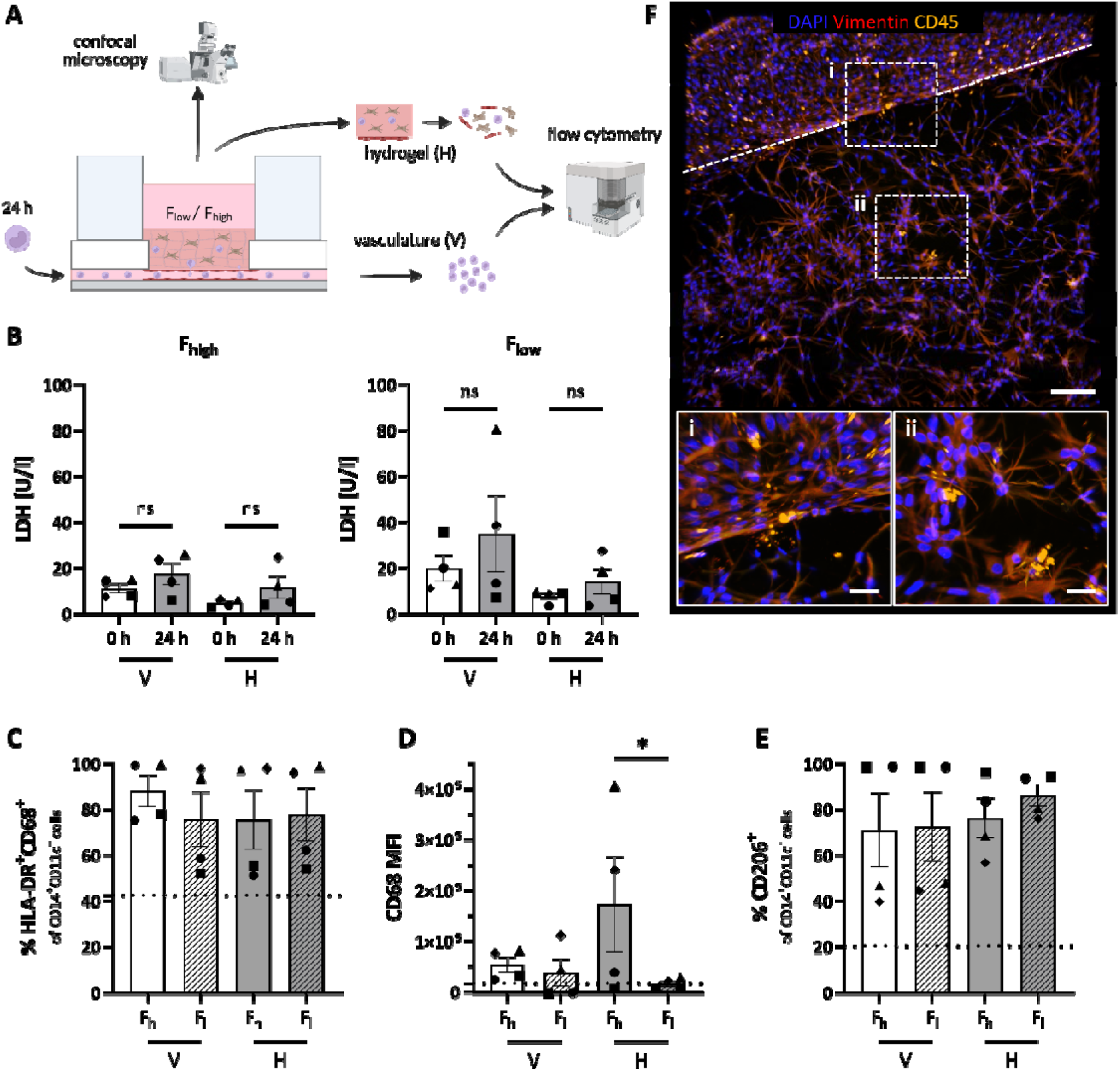
Monocyte circulation through vessel. **(A)** Monocytes recirculated through the chip vessel for 24 hours. Subsequently, cells were harvested from the vasculature (V) and the hydrogel (H) for flow cytometry analysis. **(B)** LDH analysis in the supernatant of vasculature and hydrogel before addition of the monocytes (0 h) and after 24 h circulation. **(C)** Percentage of HLA-DR^+^CD68^+^ cells within the CD14^+^CD11c^+^ cell population. **(D)** MFI of CD68 within the HLA-DR^+^CD68^+^ cells. **(E)** Percentage of CD206^+^ cells within the CD14^+^CD11c^+^ cell population. **(F)** Immune cells (CD45, yellow) attach to the vessel walls (i) and migrate into the hydrogel (ii). Vessel-hydrogel interface is indicated by a dotted line. Shown is a representative image of a F_high_ hydrogel as an example. Dotted line represents marker expression of a static control. F_h_: high fibrinogen, F_l_: low fibrinogen hydrogel concentrations, V: vasculature, H: hydrogel, LDH: lactate dehydrogenase, MFI: mean fluorescent intensity. Scale bars: 200 µm and 50 µm in magnifications. Symbols represent different donors and columns represent mean ± SEM; n = 4 independent experiments, tested with one-way ANOVA, *p ≤ 0.05, ns: not significant.

## 3. Discussion

In human *in vitro* models, vasculature is essential to overcome diffusion limits to deliver nutrients and remove waste products. In addition, it is crucially involved in the inflammatory and immune response through leukocyte transmigration.^5^

Here, we describe a method to generate a vascularized, perfused hydrogel with fibroblasts into a MOC platform and incorporate circulating peripheral blood-derived monocytes. A patterning method was used by printing a water-dissolvable sacrificial structure and connect the resulting hydrogel channel with the circulation of the commercially available HUMIMIC Chip2 24-well platform. The versatility of the method was demonstrated by variation of the cell composition as well as hydrogel component concentrations; F_low_ hydrogel to mimic a healthy tissue and F_high_ to mimic early granulation tissue. The platform is designed for maximum flexibility with different hydrogel components and cell types. Furthermore, 3D printing allows the generation of different vessel shapes by adjusting the sacrificial structure. This allows to specifically address various research questions associated with health and disease.

Perfused vessels can be generated within three days, solely by addition of an ECM and ECs. Flow is generated by a directly integrated micropump consisting of three periodically actuated pumping chambers. Pressure, frequency and direction can be modified through connection with a control unit.^27^ The pulsed flow profile generated in the HUMIMIC Chip2 96-well with empty cell culture compartments has been described earlier.^26^ We reproduced the distinct flow profile of one pumping cycle (consisting of three phases) before, within and after the hydrogel in the HUMIMIC Chip2 24-well. Notably, all three relative flow profiles were comparable with two peaks and a backflow. However, the peaks in the profile measured before the hydrogel channel were less pronounced, which may possibly be caused by the air on top of the medium in the vascular compartment acting as a buffer. A WSS of 1.7 ± 0.018 dyn/cm^2^ resembles the physiological value of small veins^28^, which is also in line with the cross-sectional dimensions of the channel (width: 500 µm, height: 100 µm). Compared to a HUMIMIC Chip2 96-well without a sacrificial structure (ca. 5 dyn/cm^2^ at 0.5 Hz)^26^, the WSS reduction is most likely caused by calculation differences and the hydrogel presence, resulting in a longer channel.

On rare occasions, detachment of the hydrogel from the hydrogel compartment or the formation of bypasses between the hydrogel and the glass bottom was observed, which can cause turbulences. Therefore, rigorous microscopic quality-control is important throughout the channel formation process, for future endothelial barrier studies. This could theoretically be solved by a coating of the chip; however, the presence of the sacrificial structure prohibits liquid coating of the hydrogel compartment after assembly as the structure would be dissolved immediately. Hence, possibilities to anchor the hydrogel to the glass bottom and the compartment walls are limited to pre-coating of individual components before assembly, which complicates the production procedure. Another limitation of our study is the PVA printing resolution, which is currently limited to 400 µm with standard extrusion printers resulting in vessel widths of 500 - 600 µm after addition of a hydrogel. Physiologically, this resembles the diameter of small arteries and veins.^29^ Even though printing at higher resolutions might be technically possible, it is very unlikely to generate capillary-like structures below 100 µm with the method described here. Accordingly, alternative techniques or combinations with these such as light-assisted printing or self-assembling of the vascular networks may be required. Compared to the initial sacrificial structure, channel width increased after addition of a hydrogel. This significant increase was slightly more pronounced in F_high_ compared to F_low_ hydrogels. Possible explanations therefore could be the contractile properties of collagen or the lower stiffness of F_high_ hydrogels.

Glucose, lactate and LDH are standard cell metabolism and cell death markers and widely used in the OoC context.^30, 31^ By measuring these markers in the medium supernatant of the perfused cultures, we show that the cells were metabolically active and viable throughout 7 days. Most importantly, for all conditions after 3 days, glucose consumption and lactate secretion reached a plateau and LDH release decreased constantly, which indicates stable (co-)culture conditions. As expected, the higher total number of cells resulted in increased glucose consumption and lactate secretion when ECs and fibroblasts were present. Also, F_high_ hydrogel conditions consumed more glucose and secreted more lactate than F_low_, indicating that cells in the F_high_ hydrogel have a higher metabolic activity than cells in F_low_ hydrogels. This is in line with the observation of vessel widening, and EC sprouting. Fibrinogen has been shown to increase sprouting of ECs into hydrogels by others as well as ourselves in the past.^32, 33^

Of note, glucose levels were higher in samples obtained from the vasculature compartment, compared to the medium control used and almost the same as levels in the hydrogel compartment. This demonstrates that glucose from the higher glucose-containing culture medium above the hydrogel diffuses through and mixes with medium in the circulation. This finding is of importance when considering the composition of culture media required to culture organoids in the different compartments within a MOC in the future. It can be assumed that if glucose diffuses through the hydrogel barrier, other components, such as serum and growth factors, will also diffuse through and mix. For day 7, we showed all 72 individual replicates across 4 repeats and 4 biological conditions to illustrate robustness of the cultures. Between different repeats, variability of glucose and lactate levels was low in the absence of fibroblasts. As expected, intra-experimental variability increased when fibroblasts were added to the hydrogel. Interestingly, glucose concentrations were more suitable to detect abnormal intra-experimental replicates. This is most likely because lactate is produced further downstream in the cell metabolism as it can be derived from pyruvate, the end product of glycolysis.^35^ For LDH release, inter-experimental variability was slightly higher and could be related to the initial seeding of EC into the microfluidics. The peak in LDH levels is a consequence of non-adherent cells dying and releasing LDH prior to their removal upon medium refreshment.

IL-6 and IL-8 are known to be secreted by both fibroblasts and ECs.^36, 37^ Chronic inflammation often precedes fibrosis, a scarring process with excess ECM secretion as a result of accumulation, proliferation, and activation of fibroblasts that eventually transition into myofibroblasts.^38, 39^ Following up on morphological changes of fibroblasts in F_high_ hydrogels which was mentioned earlier, we measured no differences in IL-6 and IL-8 levels when comparing the different hydrogel conditions. However, cytokine concentrations of TNFα, VEGF, angiopoietin-1 and PlGF were increased in the hydrogel compartment for F_high_ conditions without fibroblasts. In F_high_ hydrogels, the increased level of TNFα is an early indication of an inflamed tissue phenotype. An increased secretion of the well characterized angiogenic factors VEGF, angiopoietin-1 and PlGF by ECs could have caused the observed angiogenic sprouting.^40^ Surprisingly, no difference in cytokine secretion between F_high_ and F_low_ hydrogels was observed when fibroblasts were present. An explanation therefore could be that fibroblasts regulate EC cytokine secretion or take up cytokines which results in lower cytokine concentrations in the supernatant.

As a first step towards an immune-competent model, we made use of the perfusability of the blood vessel and added monocytes into the circulation. The CD14^+^ monocytes applied to the chip differentiated into HLA-DR^+^CD68^+^ macrophage-like cells under flow. Notably, the cells became more macrophage-like in F_high_ compared to F_low_ hydrogels, as a high CD68 MFI indicates a large quantity of the receptor and activated macrophages.^41^ Further characterization of the cells for expression of CD40 (M1) and CD206 (M2) macrophages demonstrated that an M2-like phenotype was acquired on-chip. These macrophages are typically regarded as anti-inflammatory and involved in the processes of wound healing and tissue repair.^42^ In adipose tissue, CD206^+^ tissue resident macrophages have been found to contribute to tissue homeostasis.^43^ However, upregulation of this marker in both hydrogel conditions suggests either an effect of flow or paracrine signaling from the other cell types leading to this differentiation. The variability in the degree of macrophage differentiation, CD68 MFI and percentage of CD206 expression within the experimental conditions can most likely be attributed to donor variation (different fibroblast and EC donors were used in each experiment) as large donor variation has previously been reported for ECs in particular.^33^ Other studies have shown that ECs can interact with macrophages, influence their phenotype, and even differentiate them into an M2-like phenotype.^44^ Of note, even though an additional washing step was added before digesting the hydrogels for flow cytometry analysis, it was not possible to distinguish between immune cells that have transmigrated into the hydrogel and those that have adhered to the ECs lining the channels. We therefore also visualized the transmigrated cells in the hydrogel by confocal microscopy. Migration of CD45^+^ cells into the hydrogel was confirmed, which shows the cells’ ability to trans-migrate across the endothelium into the tissue and confirms the use of the model for future immune studies.

In future experiments, activation and migration of immune cells could be triggered, e.g., by the addition of pro-inflammatory cytokines such as TNFα, lipopolysaccharide, and living or heat-killed bacteria. The open top design of the chip enables the generation of barrier models (e.g., lung, intestine or skin) above the vascularized hydrogel, which would allow apical bacterial exposure and the study of host-microbiome immune responses. The MOC platform allows the use of additional immune-competent organ models such as the, lymph node and other secondary lymphoid organs.^45^ In future studies, the method presented here can be combined with the HUMIMIC Chip4 to connect up to five different vascularized organ models on a single MOC.

## 4. Conclusions

There is an unprecedented need to create more physiological tissue and organ models, not only in research to understand human (patho-)physiology, but also in toxicology, drug discovery, and in regenerative and personalized medicine. In the past years, tremendous progress has been made in the fields of MPS, 3D (bio-)printing, ECM, and combinations of these, making it possible to increase complexity of organ models by vascularization.^46, 47^ However, these models often rely on perfusion of external tubing and pumps to generate perfusion and require complex experimental setups, making them extremely difficult to reproduce.

Our study presents a versatile and easy method to generate vascularized perfused hydrogels in a MOC platform. Stable blood vessels can be generated within three days and hydrogel concentrations can be modified to generate a healthy or diseased vessel. In high fibrinogen hydrogels, angiogenesis, increased secretion of angiogenic and inflammatory cytokine secretion and presence of anti-inflammatory macrophages are hallmarks of the early phase in wound healing and early granulation tissue formation.^34^ Therefore, we demonstrate that simple adjustments to the hydrogel composition can impact cellular metabolism and behavior, indicating early granulation tissue. This makes it a valuable tool to engineer the next generation of vascularized and immune-competent organ models in the future.

## 5. Methods

### Cell culture

Human skin from healthy donors was collected as surgical waste after abdominal dermolipectomy. The tissue was used in an anonymous fashion and all donors gave informed consent as described in the ‘Code of Conduct for Health Research’ as formulated by COREON (Committee on Regulation of Health Research; https://www.coreon.org) and the following procedures were approved by the institutional review board of the Amsterdam UMC.

### Fibroblasts and endothelial cells

Dermal fibroblasts were isolated from human skin as described earlier.^48^ Cells were cultured in fibroblast medium consisting of Dulbecco’s modified Eagle’s medium (DMEM; Thermo Fisher Scientific, Waltham, MA, USA), supplemented with 1% UltroserG (BioSepra, Cergy, France) and 1% penicillin/streptomycin (P/S; Invitrogen, Paisley, UK). Cells were cultured at 37 °C, 5% CO_2_, medium was exchanged every 3-4 days and cells were used at passage ≤ 3. Dermal endothelial cells (ECs) were isolated from human dermis. After enzymatic digestion of the dermal fraction of the skin, cells were cultured in Endothelial Cell Growth Medium MV2 (MV2; PromoCell, Heidelberg, Germany), always supplemented with 1% P/S. When endothelial colonies were formed, the endothelial population was enriched with a CD31 MicroBead Kit (Miltenyi Biotec, Bergisch Gladbach, Germany). Cells were amplified in MV2 and after harvesting, ECs were labeled with anti-human podoplanin Alexa-488 and anti-human CD31 PE (both BioLegend, San Diego, CA, USA) and subsequently sorted on a BD FACSAria™ Fusion Flow Cytometer (BD Biosciences, Franklin Lakes, NJ, USA) into a pure population of CD31^+^PDPN^-^ blood endothelial cells (BECs). Cells were cultured at 37 °C, 5% CO_2_ in MV2 and medium was exchanged every 3-4 days. Cells at passage ≤ 8 were used for experiments.

### Peripheral blood derived monocytes

Human monocytes were isolated using magnetic-activated cell sorting with CD14^+^ magnetic beads from peripheral blood mononuclear cells (PBMCs) as previously described.^49^ At day 3, monocytes were added to the MOC for 24 hours at 3.5 x 10^5^ cells per chip in media composed of RPMI-1640 (Lonza, Basel, Switzerland) supplemented with 10% fetal calf serum (FCS; GE Healthcare, Chicago, IL, USA), 50 µM β-mercaptoethanol (Merck, Darmstadt, Germany), 100 IU/ml sodium-penicillin (Thermo Fisher Scientific), 100 µg/ml streptomycin (Thermo Fisher Scientific), and 2 mM L-glutamine (Thermo Fisher Scientific).

### Microphysiological system (multi-organ-chip)

A HUMIMIC Chip2 24-well (TissUse) was used with a sacrificial print, aligned with the microfluidic channels of the chip (Supplementary Figure S1A). The computer aided design (CAD) model to generate the PVA structure was created in solidworks (v2018; Dassault Systèmes SOLIDWORKS, Waltham, MA, USA) and printed with an Ultimaker S5 (Ultimaker, Utrecht, The Netherlands); BB print core and a 0.4 mm nozzle, according to the manufacturer’s instructions. Recirculating Flow was applied with a HUMUMIC Starter (TissUse) control unit at 0.5 Hz, ± 500 mbar, counterclockwise.

### Hydrogels

Two different hydrogels containing differing ratios of collagen and fibrinogen were used in experiments, based on previous publications. The concentration for the F_low_ hydrogels was optimized for reconstructed human skin models based on the minimum concentration of fibrinogen to prevent shrinkage.^50^ F_high_ concentrations were optimized and selected for EC sprouting and microvasculature formation on-chip.^51, 52^ Collagen was isolated from rat tails and dissolved in 0.1% acetic acid (VWR, Radnor, PA). Then, 10x HBSS (Thermo Fisher Scientific) was added to the collagen solution (diluted 1:8) and the pH neutralized with 1M NaOH. Low fibrinogen (F_low_) hydrogels were composed of a final concentration of 3 mg/ml collagen and 1 mg/ml fibrinogen (Diagnostica Stago, Paris, France). For high fibrinogen (F_high_) conditions, hydrogels were composed of a final concentration of 2.5 mg/ml fibrinogen and 0.1 mg/ml collagen. To the collagen/fibrinogen mixture of both hydrogel conditions, aprotinin (SERVA, Heidelberg, Germany) and thrombin (Merck) were added at final concentrations of 50 KIU/ml and 0.5 U/ml respectively and resuspended thoroughly before casting. 350 µl hydrogel with or without 7 x 10^4^ fibroblasts/ml was casted into the hydrogel compartment of the chip, covering the sacrificial PVA structure. Chips with hydrogels were subsequently incubated at 37 °C for 30 min for collagen-fibrin network formation and polymerization. Next, 300 µl MV2 was added to the vascular compartment and 500 µl fibroblast medium to the hydrogel compartment and the chip was connected to the HUMIMIC Starter to apply flow, then incubated overnight at 37 °C, 5% CO_2_.

### Blood endothelial cell seeding

To flush out residual printed sacrificial structures, the vascular compartment was washed with 300 µl MV2 before seeding the cells. Then, 5 x 10^5^ BECs were seeded into the patterned hydrogel structure via the pressure differences after adding the cell suspension to the vascular compartment. Chips were subsequently incubated at 37 °C, 5% CO_2_ for 4 hours. After 4 hours, the vascular compartment was washed with 300 µl MV2. Then, 300 µl fresh MV2 was added to the vascular compartment and 500 µl fibroblast medium on top of the hydrogel, both media supplemented with 50 KIU/ml aprotinin. MV2 and fibroblast medium were always supplemented with 50 KIU/ml aprotinin from here on. A full medium exchange was performed on day 3 and day 5 for both compartments: in the vascular compartment 300 µl MV2 was collected and replaced with fresh medium and for the hydrogel compartment, 500 µl fibroblast medium was collected and replaced with fresh medium. At day 7, medium from the different compartments was collected. For all time points, media was stored separately at 4 °C until metabolic analysis was performed. Afterwards, media was stored at -20 °C until further analysis. As a quality-control, bright field images were taken at day 7 to ensure no major detachment of the hydrogel from the cell culture compartment (Supplementary Figure S5).

### Fluid dynamics characterization - flow rates and shear stress

To characterize the flow dynamics, the flow velocity within the channels was measured with time and spatial resolution using micro-particle image velocimetry (µPIV). The measurement method was described previously in literature.^53^ First, chips were filled with 0.4% w/v 5 µm Aldehyde/Sulfate Latex Beads in PBS (Thermo Fisher Scientific) as tracer particles and 0.1% Triton-X-100 (VWR) to reduce the sedimentation. The chips were then connected to a HUMIMIC Starter (TissUse) and perfused with a frequency of 0.5 Hz, and pressure/vacuum of ± 500 mbar. The µPIV measurements were performed using an inverted microscope Axio Vert.A1 (Zeiss, Jena, Germany) with a 2.5x objective, LED light source and a high-speed camera HXC40 (Baumer, Friedberg, Germany). For the measurement, the center of the channel was focused to capture the maximum velocity of the particles. The recorded image series were analyzed with the open source toolbox PIVLab for MATLAB for mean flow velocity and its extrema.^54^ Flow rates were normalized with a min-max normalization.

For a rectangular channel, WSS was calculated according to Hagen-Poiseuille’s law from the dynamic viscosity η of the medium, the mean flow velocity Q and the width w and height h of the channel (adapted from ^55^):

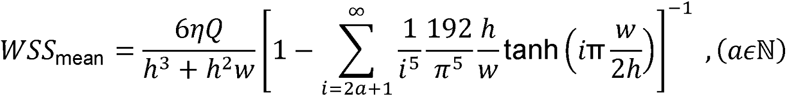

As it was not possible to determine the WSS for channels with complex cross-sectional geometries analytically, the WSS for the hydrogel channel was simulated using COMSOL Multiphysics 5.2a (COMSOL, Stockholm, Sweden). For this purpose, the cross-sectional geometry of the channels was measured with a Nikon AX R confocal microscope (Nikon, Tokyo, Japan) and modulated three-dimensionally as a straight channel. The simulation was carried out as a 3D single-phase steady fluid flow. It was assumed that the flow is laminar. Derived from the continuity equation, it can be assumed that the mean flow in the hydrogel channel is equal to the mean flow in the PDMS channels. Consequently, the previously measured mean volume flow through the hydrogel channel was used for the simulation.

### Immunofluorescent and histological staining

All flow incubations conducted in this section were performed with the following perfusion settings: 0.5 Hz, ± 500 mbar, counterclockwise. Hydrogels were preserved in chips directly by adding 4% paraformaldehyde (Electron Microscopy Sciences, Hatfield, PA, USA), followed by incubation for 10 min in flow and 1 hour static at RT. Chips were then pre-treated with 0.2% Triton in PBS, 10 minutes in flow at RT before the immunostaining. Next, chips (100 µl to the vascular and 200 µm to the hydrogel compartment on top of the hydrogel) were incubated with mouse anti-human CD31 (Dako, Glostrup, Denmark), incubated for 30 minutes in flow at RT and afterwards overnight static at 4 °C. The following day, chips were washed twice with 0.2% Triton in PBS and then incubated with goat anti-mouse 488 (Thermo Fisher Scientific), directly labeled anti-vimentin Alexa Fluor 647 (BioLegend) and DAPI (Invitrogen) for 30 minutes in flow at RT followed by an overnight static incubation at 4 °C. The following day, chips were washed twice with 0.2% Triton in PBS and stored at 4 °C until analysis. For histological stainings, hydrogels were removed from the chips and fixed overnight in 4% formaldehyde (VWR). Then, samples were embedded in paraffin and 5 µm sections were used for a hematoxylin & eosin (H&E) staining to investigate hydrogel degradation (Supplementary Figure S4F).

### FITC-dextran permeation analysis

FITC-dextran (70 kDA; Sigma-Aldrich, St. Louis, MO, USA) was dissolved in PBS and used in a final concentration of 0.5 mg/ml. 100 µl were added to the vascular compartment of the MOC and images were taken after a 10 minute perfusion.

### Microscopy and channel width measurements

Bright field images and FITC-dextran perfusion images were taken with a Keyence BZ-X700E (Keyence, Osaka, Japan) and analyzed in the BZ-X800 Analyzer (v1.1.2; Keyence). Channel widths were measured in bright field, perpendicular to the channels in triplicates for ≥ 11 individual circulations per condition, either 24 hours after the gel was casted or with cells at day 7 (Supplementary Figure S6). Fluorescent images were taken with a Nikon ECLIPSE Ti2 (Nikon, Tokyo, Japan) and confocal images with a Nikon AX R (Nikon). Images were analyzed in Imaris (v10.1.0; Oxford Instruments, Oxfordshire, UK).

### Viability and metabolic analysis

To measure levels of glucose, lactate and lactate dehydrogenase (LDH) activity in the absence of monocytes (performed at TissUse, Berlin), medium was analyzed on day 0 (before addition of ECs), 3, 5 and 7, with an Indiko™ Plus Clinical Chemistry Analyzer (Thermo Fisher Scientific) in combination with the Glucose (HK) 981779 kit (Thermo Fisher Scientific), Lactate Fluitest LA 3011 kit (Analyticon Biotechnologies, Lichtenfels, Germany), and LDH IFCC 981782 kit (Thermo Fisher Scientific). For experiments with monocytes (performed at Amsterdam UMC), the Cytotoxicity Detection Kit PLUS (LDH) was used (Figure 7; Roche, Basel, Switzerland), according to the manufacturer’s instructions. MV2 and fibroblast medium were used as controls and basal glucose levels in the whole circulation calculated with the ratio of 3/8 vasculature and 5/8 hydrogel medium. For lactate and LDH analysis, basal medium levels were subtracted and set to 0 when below medium levels.

### Cytokine analysis

The Human Angiogenesis Panel V02 (BioLegend, San Diego, CA, USA) cytokine bead array was used to measure concentrations of Interleukin-6 (IL-6), IL-8, tumor necrosis factor α (TNFα), basic fibroblast growth factor (bFGF), vascular endothelial growth factor (VEGF), soluble platelet endothelial cell adhesion molecule (sPECAM-1), epidermal growth factor (EGF), placental growth factor (PlGF), Angiopoeitin-1 and Angiopoeitin-2. The assay was performed according to the manufacturer’s instructions and acquired on an Attune NxT flow cytometer (Thermo Fisher Scientific). Data was analyzed with the LEGENDplex Data Analysis Software Suite (BioLegend).

### Multispectral flow cytometry

Cells were added to the vascular compartment at day 3 and circulated on the chip for 24 hours (Supplementary Figure S4A). Then, cells were collected from the vasculature compartment. The vasculature compartment was washed with fresh medium by pumping at RT for 10 min to remove remaining cells which did not attach. Hydrogels were subsequently enzymatically digested into a single cell suspension for flow cytometry. Hydrogels were incubated in 0.2 mg/ml Collagenase P and 0.1 mg/ml DNase I (both from Sigma-Aldrich) for 1 h at 37 °C. Afterwards, the reaction was stopped with 2% FCS and 5 mM EDTA (Sigma-Aldrich) in PBS.

Cell suspensions were stained in a 96-well plate at 4 °C for multispectral flow cytometry analysis. Cells were first washed with PBS and stained with a fixable viability dye in 1:2000 (eFluor™ 780 labeled; Invitrogen) for 10 minutes at 4 °C. Next, cells were washed with 0.1% BSA and 0.05% NaN_3_ in PBS (FACS buffer) prior to Fc-receptor blocking with 10% normal human serum in FACS buffer. Cells were then incubated with directly labeled antibodies, against extracellular markers, diluted in blocking buffer in the following dilutions: anti-CD11c in 1:200 (BUV661 labeled), anti-CD14 in 1:50 (pacific blue labeled), anti-HLA-DR in 1:100 (BV711 labeled), anti-CD206 in 1:50 (FITC labeled) and anti-CD40 in 1:50 (APC labeled; all BioLegend). After staining, cells were washed two times with FACS buffer and fixed with 2% PFA (VWR) for 10 minutes at RT. Cells were then washed with 0.5% Saponin buffer, and the intracellular antibody was added, anti-CD68 in 1:100 (BV785 labeled; BioLegend). Samples and single stains on beads were acquired on Aurora 5-laser Flow Cytometer (Cytek; Fremont, CA, USA). Data was analyzed with FlowJo (v10.7; Ashland, OR, USA) and OMIQ (Boston, MA, USA). Analysis of monocytes and macrophages was based on a gating strategy to exclude other cell types such as ECs and fibroblasts (Supplementary Figure S4B). Monocytes were gated on cells, single cells, live cells, CD14^+^CD11c^+^HLADR^+^CD68^-^ and macrophages on CD14^+^CD11c^+^HLADR^+^CD68^+^ (Supplementary Figure S4C).

*Rheology analysis:* To measure the hydrogel stiffness, F_high_ or F_low_ hydrogels were constructed with fibroblasts and ECs in the chips as described earlier and analyzed at day 4, either with or without monocytes (Figure 4D and Supplementary Figure S4E). Hydrogels were measured directly in the chips with a Chiaro nanoindenter (Optics 11 life, Amsterdam, The Netherlands). For indentation, a probe with 0.025 N/m stiffness and 25 µm tip radius was used in the peak load poking mode, and a maximum load of 0.02 µN. Shortest distance between two measurements was 200 µm.

### Statistical analysis

Data is presented as mean ± standard error of the mean. The number of different skin donors and the number of independent experiments is described in the legends of each figure. GraphPad Prism (v9.5.1; GraphPad Software Inc., La Jolla, CA, USA) was used for statistical analysis with tests indicated in figure legends. Differences were considered as significant when p ≤ 0.05.

## Supporting Information

Supporting Information is available from the Wiley Online Library or from the author.

## Author Contributions

Conceptualization: JJ, ED, SG, JJK; Funding Acquisition: SG; Investigation: JJ, AIM, HE, MT; Methodology: JJ, PB, AIM, ED, SG, JJK; Project administration: ED, SG, JJK; Visualization: JJ, HE; Writing-Original Draft Preparation: JJ, SG, JJK; Writing-Review and Editing: JJ, AIM, HE, ED, SG, JJK.

## Acknowledgments

The presented work was funded by the SMART Organ-on-Chip consortium, NWO-TTW Perspective Programme of the Dutch Research Council (NWO; project no. P19-03), the European Union’s Horizon 2020 research and innovation program (grant agreement no. 847551) ARCAID, the LymphChip consortium, NWA-ORC programme of the Dutch Research Council (NWO; project no. 1292.19.019), the ReSHAPE consortium, EU-H2020 (grant agreement no. 825392), and the BIRDIE consortium, EU-H2020 (grant agreement no. 964452). Views and opinions expressed are however those of the author(s) only and do not necessarily reflect those of the European Union or the European Health and Digital Executive Agency (HADEA). Neither the European Union nor the granting authority can be held responsible for them. The authors would like to thank Linda Henriette Reichel for PIV measurements and analysis, Artur Christian Garcia da Silva for the FITC-dextran permeation assay and the Microscopy & Cytometry Core Facility at Amsterdam UMC (location Vrije Universiteit Amsterdam) for their support.

## Supporting Information

**Figure S1: Experimental timeline and fit of the PVA structure**. Additional information for Figure 1. **(A)** Bright-field image of the PVA-PDMS interface shows fit of the sacrificial structure in the PDMS channel. Scale bar: 1000 µm. **(B)** Chips were produced and hydrogel (+/- fibroblasts) and ECs added. Subsequently, the vessels were perfused for 7 days. ECs: endothelial cells, ME: medium exchange, LDH: lactate dehydrogenase, F_high_: high fibrinogen, F_low_: low fibrinogen hydrogel concentrations.

**Figure S2: Metabolic markers of EC-fibroblast co-cultures show low variability between different repeats.** Individual repeats of Figure 6. Variability between different repeats and inter-experimental variability of a total of n = 72 (_≙_ 36 chips) individual measurements at day 7. For vasculature and hydrogel measurements of glucose, lactate and LDH, four conditions are shown with intra-experimental replicates (each column) and repeats (1-4). Dotted lines indicate basal glucose levels of respective vasculature and hydrogel medium and dashed lines the basal glucose levels in the whole chip. Symbols represent different conditions and line mean of intra-experimental replicates.

**Figure S3: Variability of cytokine secretion.** Individual repeats of Figure 5. Measurement of 9 different cytokines shows variability between different repeats and inter-experimental variability of a total of N = 72 (≙ 36 chips). Shown are individual measurements at day 7. Black data points were outside the detection limit, extrapolated in this figure and set to detection limit in Figure 5.

**Figure S4: Experimental timeline, gating strategy, histological sections and stiffness measurements for monocytes and macrophages circulated through the vessel-on-chip. Addition to Figure 7. (A)** Experimental timeline for 24 h monocyte perfusion on-chip. **(B)** Monocytes were directly gated based on size and granularity (FSC-SSC) to initially remove other cell types such as fibroblasts and ECs together with debris. Hydrogel and vasculature sample of the same circulation is shown as an example. **(C)** Gating was set on cells, single cells, live cells, CD14^+^CD11c^+^ cells, HLADR^+^CD68^+^ and CD40^+^ or CD206^+^ for M1 and M2 macrophages. FSC: forwards scatter, SSC: side scatter. **(D)** Percentage of CD40^+^ cells within the CD14^+^CD11c^+^ cell population. Dotted line represents marker expression of a static control. F_h_: high fibrinogen, F_l_: low fibrinogen hydrogel concentration, V: vasculature, H: hydrogel. Symbols represent different donors and columns represent mean ± SEM; n = 4 independent experiments. **(E)** Stiffness of different hydrogel conditions, measured after 4 days in the chip with fibroblasts and ECs and 24 hours of monocyte perfusion. Each condition with n ≥ 20 data points. Unpaired t-test, ****p < 0.0001. **(F)** Cross-sectional H&E staining of F_high_ and F_low_ hydrogels with fibroblasts, ECs, and monocytes. The perfusable endothelialized channel is shown in the center (*). Images were taken at day 4 after a 24 hour perfusion of monocytes. H&E: hematoxylin & eosin. Scale bars: 100 µm.

**Figure S5: Hydrogel characterization and quality-control.** Gross bright-field images of F_high_ and F_low_ hydrogels in chip with ECs and fibroblasts at day 7. Shown are representative images for 3 different replicates. Scale bars: 2000 µm.

**Figure S6: Vessel width measurements.** Additional information for Figure 4C. Addition of a hydrogel on top of the sacrificial structure induced widening of the channel. Images were taken before addition of the hydrogel and after 24 hours. Red lines indicate channel alignment and perpendicular width measurement. Scale bars: 1000 µm in overviews, 500 µm in magnifications.

**Movie S1: Hydrogel channel rendering.** ECs were stained with vimentin (yellow) and nuclei with DAPI (blue) at day 7. Surface-rendering is shown in red.

