## Supplementary Figures for "A Sacrificial 3D Printed Vessel-on-Chip Demonstrates a Versatile Approach to Model Connective Tissue Pathology"

### Supplementary Figure S1

**A**

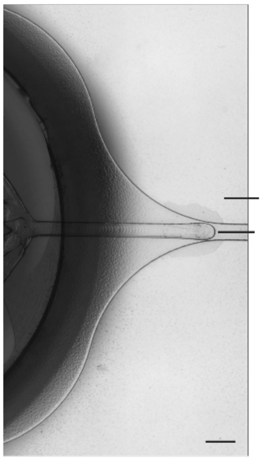


**B**

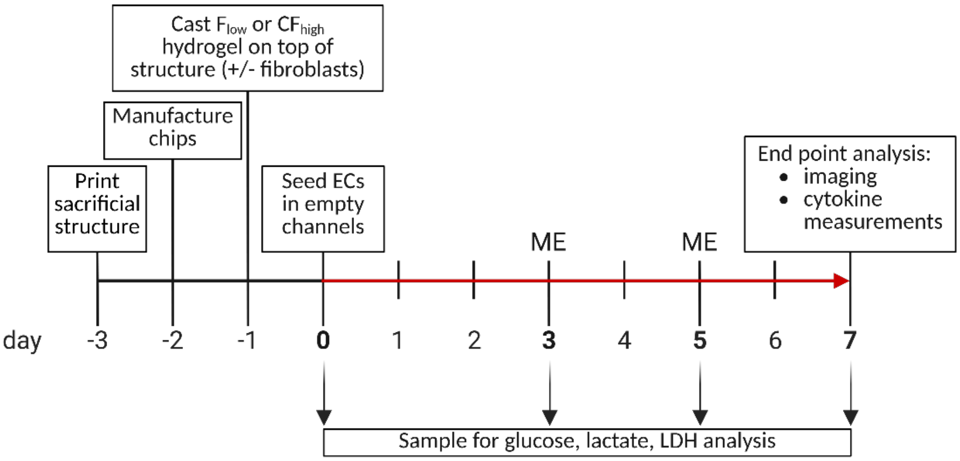

### Supplementary Figure S2

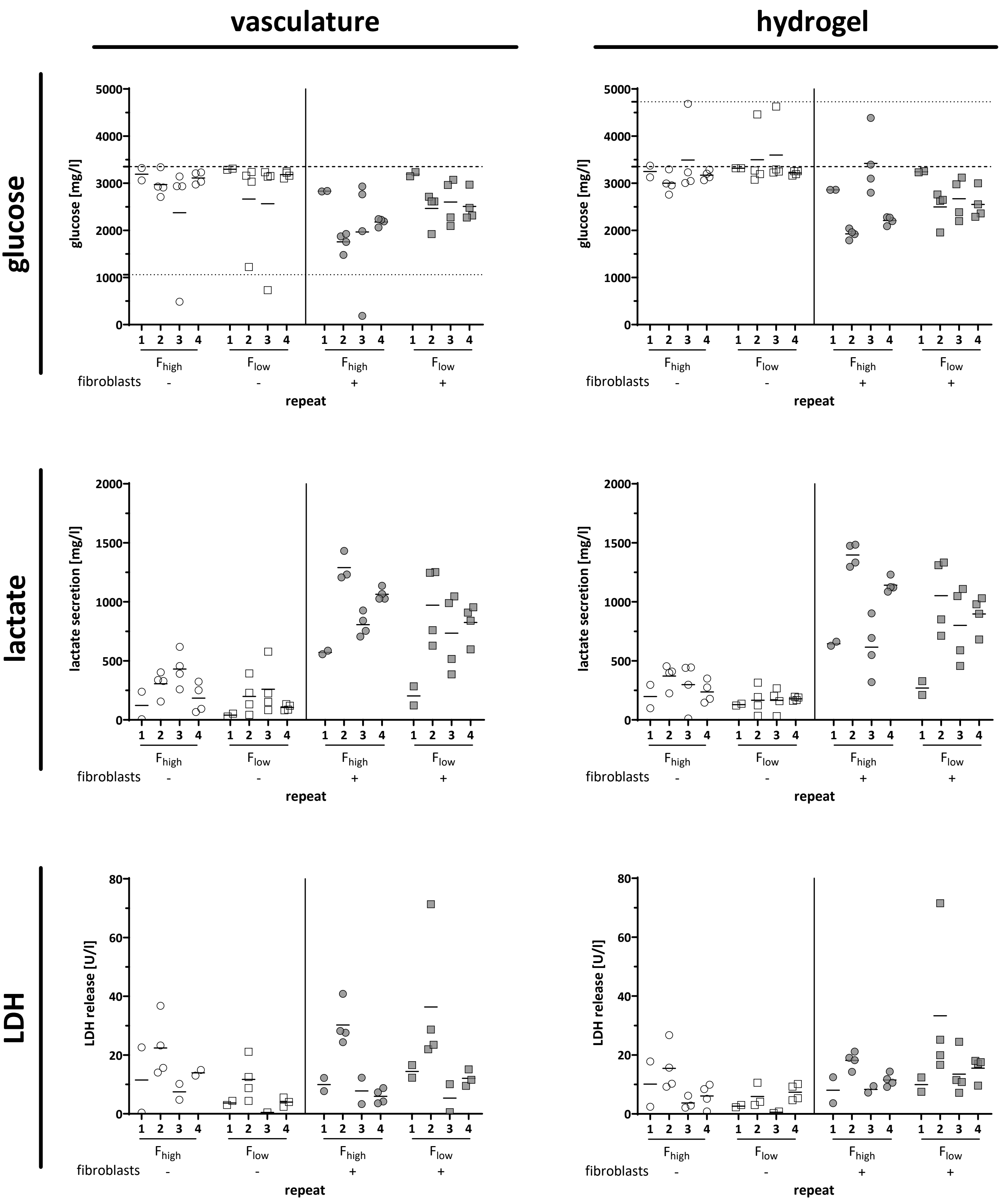

Supplementary Figure S3

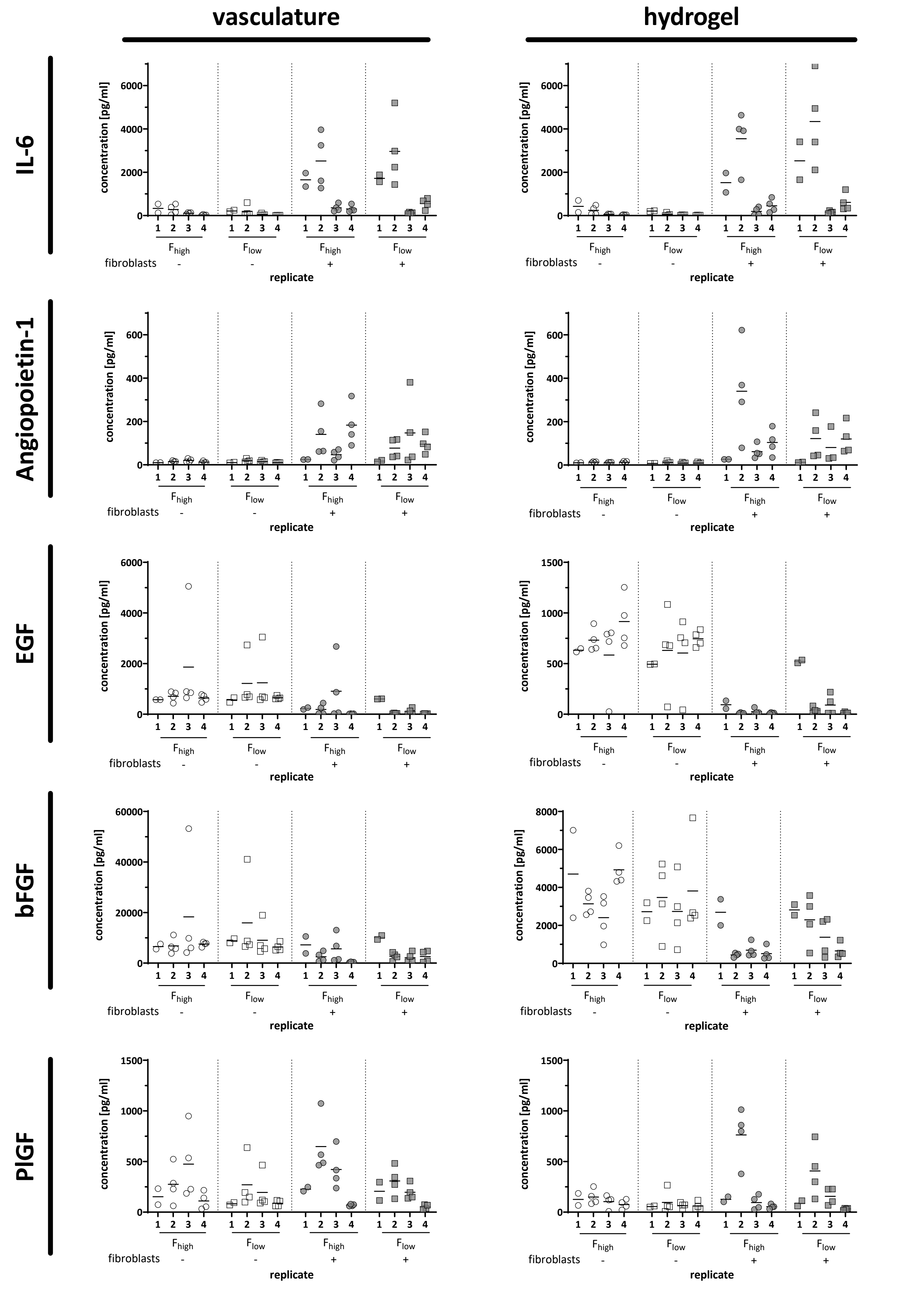

### Supplementary Figure S3, continued

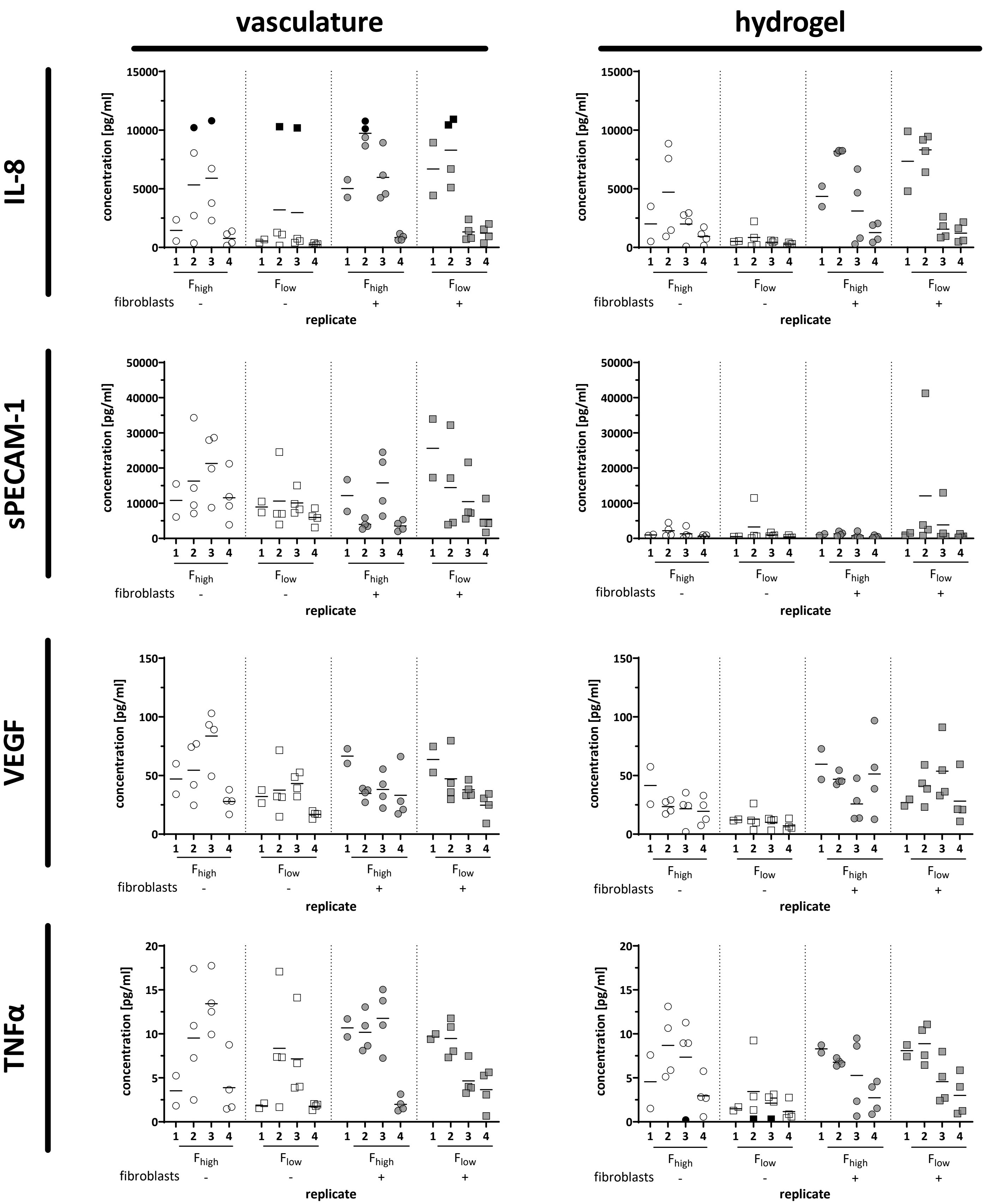

### Supplementary Figure S4

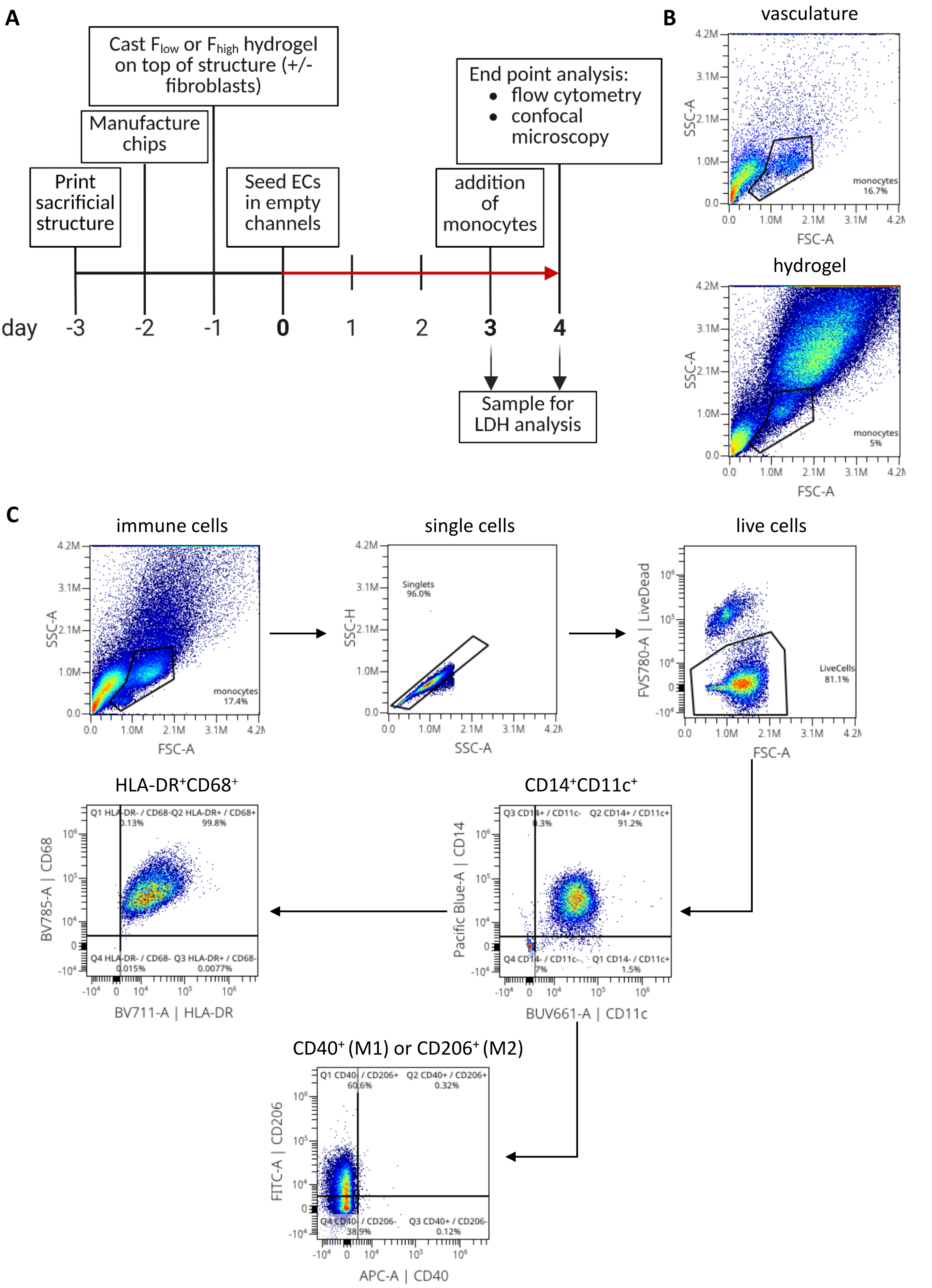

Supplementary Figure S4, continued

D

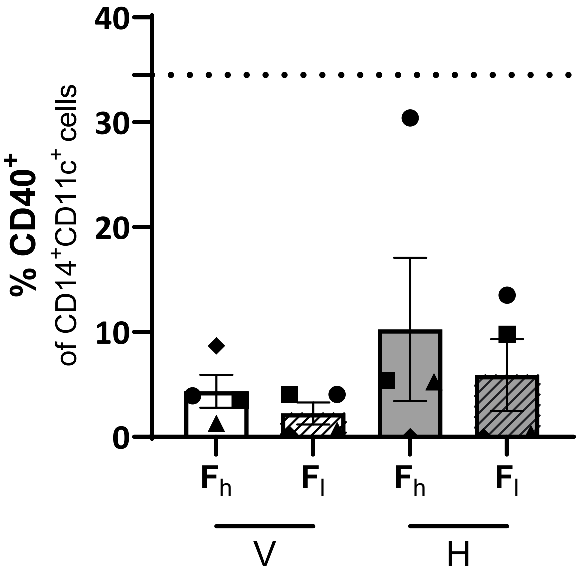


E

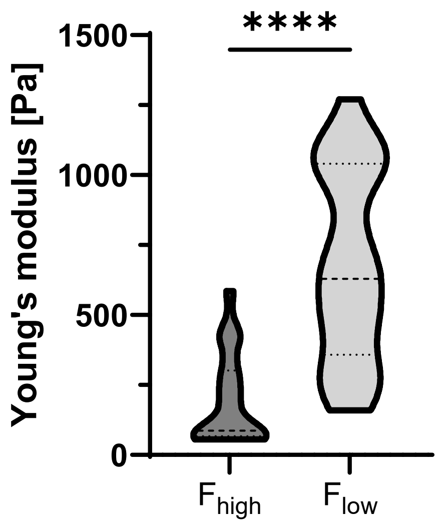


F

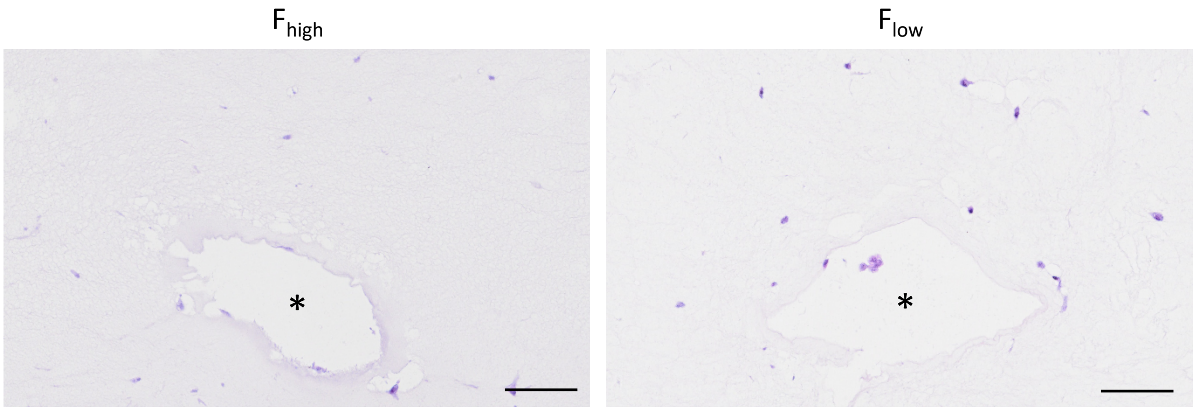

### Supplementary Figure S5

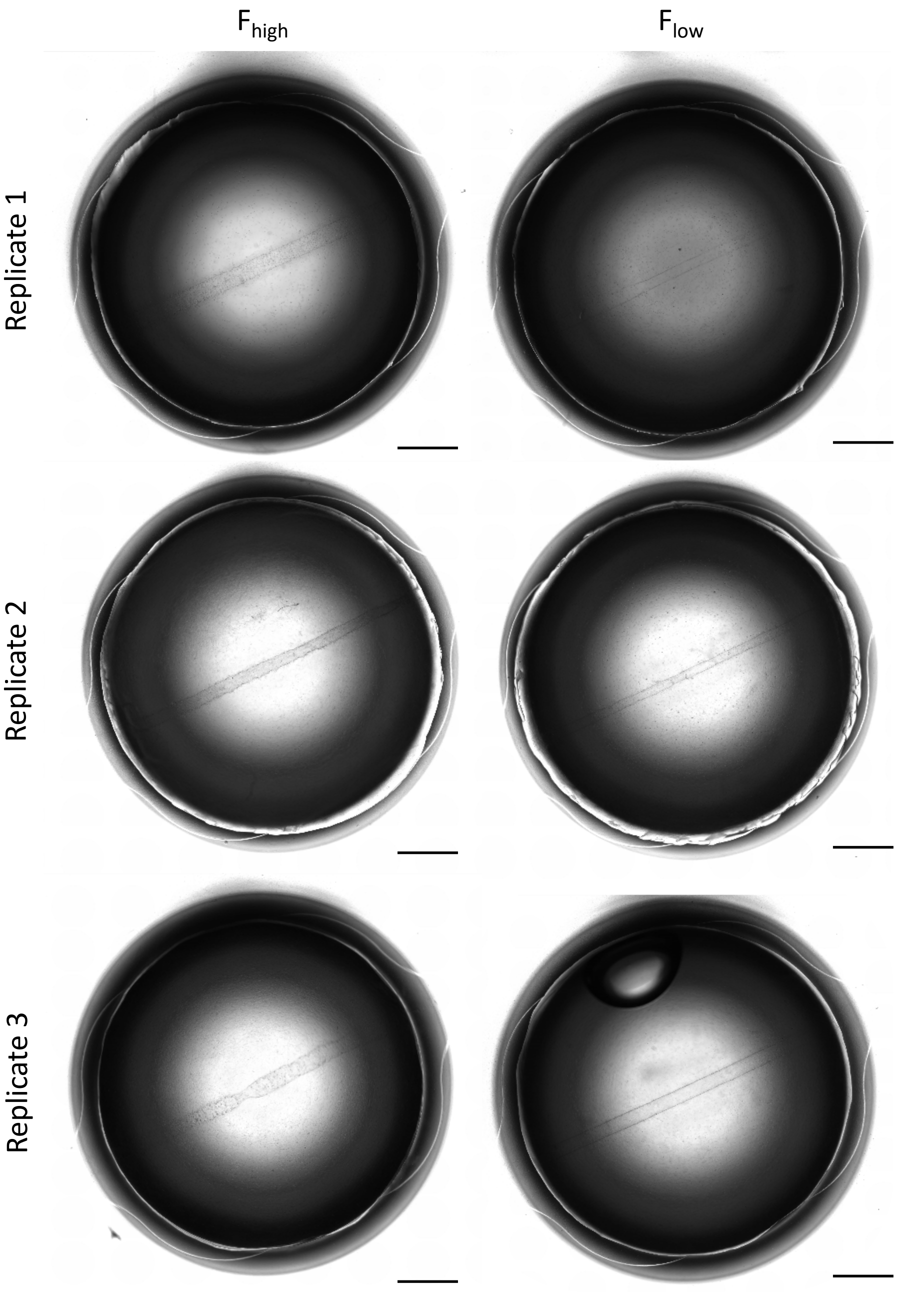

### Supplementary Figure S6

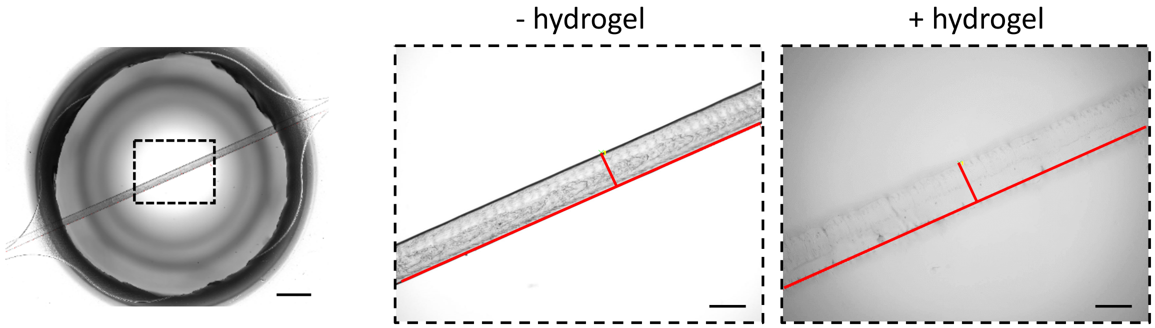
